# KIF5A binds RNA to orchestrate synaptic mRNA localization and stress granules in ALS

**DOI:** 10.1101/2025.10.31.685813

**Authors:** Phuong Le, Neeraj K. Lal, Shuhao Xu, Sara Mumford, Megan Huang, Dong Yang, Orel Mizrahi, Benjamin Hoover, Brian Yee, Yuan Mei, Katherine Rothamel, Hsuan-Lin Her, Steven M. Blue, Neil A. Shneider, Gene W. Yeo

## Abstract

Neuronal health depends on the precise transport and local translation of mRNAs to maintain synaptic function across highly polarized cellular architecture. While kinesin motor proteins are known to mediate mRNA transport, the specificity and direct involvement of individual kinesins as RNA-binding proteins (RBPs) remain unclear. Here, we demonstrate that KIF5A, a neuron-specific kinesin implicated in amyotrophic lateral sclerosis (ALS), functions as an RBP. We show that KIF5A directly binds mRNAs encoding synaptic ribosomal proteins and is required for their synaptic localization and for maintaining normal synaptic composition and function. Additionally, we show ALS-linked KIF5A mutations confer gain-of-function properties, enhancing mRNA binding, increasing synaptic ribosomal protein accumulation, inducing neuronal hyperexcitability, and impairing stress responses. These findings reveal a previously unrecognized mechanism by which mutant KIF5A disrupts synaptic homeostasis. Our work positions a kinesin motor protein as an RBP with critical roles in mRNA transport, local translation, and stress response.

**Highlights:**

- KIF5A interacts with mRNA encoding synaptic ribosomal proteins
- KIF5A is required for normal synaptic composition and function
- KIF5A binds to G3BP1 and G3BP1 stress granule associated proteins
- KIF5A mutant ALS patient-derived motor neurons have abnormal synaptic function and stress response

## INTRODUCTION

Neurons are among the most structurally complex and polarized cells in the human body, with axonal projections that can extend up to 1 meter in length^1^. This extraordinary architecture presents a significant logistical challenge: the constant activity at synaptic terminals requires a continuous and localized supply of new proteins at sites that may be far removed from the cell body. Relying solely on the transport of proteins synthesized in the soma is insufficient to meet these demands, as passive diffusion is extremely slow—estimated at approximately 0.1 μm/s^2^, which would take up to 120 days for proteins to reach distal synapses. Even active transport along microtubules by motor proteins, though faster at roughly 2 μm/s^3,4^, could still require up to six days for delivery, which is incompatible with the half-life of many proteins^5,6^ and the rapid dynamics of synaptic function.

The need for a more efficient mechanism led to the hypothesis that neurons locally translate proteins at synaptic sites, a concept first supported by electron microscopy studies in the 1970s that revealed the presence of polyribosomes within dendritic spines^7,8^. These findings suggested that mRNAs are transported into neurites, where they are locally translated to meet the immediate and spatially restricted demands of synaptic activity. However, direct evidence for the transport of mRNAs into neurites and their local translation only emerged in the early 2010s^9^, fundamentally transforming our understanding of neuronal protein synthesis and synaptic plasticity.

It is now widely accepted that mRNAs are transported in ribonucleoprotein (RNP) complexes or RNA transport granules along microtubules, mediated by motor proteins such as kinesins^10^. Despite this perspective, key questions remain unresolved: (1) which mRNAs are selectively transported, and (2) what the precise mode of interaction between mRNAs and motor proteins is—specifically, whether this occurs through direct binding or via adaptor proteins. As the list of canonical RNA-binding proteins (RBPs) expands^11–14^, the possibility that certain kinesin family members may themselves function as RBPs is an intriguing area of investigation.

Among the kinesin superfamily, KIF5A stands out due to its high and preferential expression in the brain and its established links to neurodegenerative and neurodevelopmental disorders, including amyotrophic lateral sclerosis (ALS), hereditary spastic paraplegia (SPG10), and Charcot-Marie-Tooth disease type 2 (CMT2)^15–19^. Like other kinesins, KIF5A is composed of three domains: an N-terminal motor domain, a central stalk domain, and a C-terminal cargo-binding domain. Notably, mutations associated with SPG10 and CMT2 are found in the motor domain, while ALS-associated mutations cluster in the cargo-binding domain, implicating altered cargo interactions in disease pathogenesis^15^. Previous studies have shown that ALS-linked mutations in KIF5A disrupt cargo binding, yet the mechanisms by which these alterations contribute to neurodegeneration remain unclear^20,21^.

Our study addresses these critical gaps by investigating whether KIF5A can function as an RNA-binding protein that interacts with specific mRNAs and regulate their localization at the synapse. We delineate the role of KIF5A in synaptic composition and function through a combination of genomics, biochemical, imaging and electrophysiological approaches, leveraging patient-derived stem cell models. Furthermore, we explain how ALS-associated mutations in KIF5A alter its RNA and protein interactions, leading to gain-of-function phenotypes that disrupt synaptic homeostasis and stress responses. These findings provide new mechanistic insights into the contribution of altered mRNA transport and stress pathway to ALS pathogenesis and position KIF5A as a key regulator of neuronal mRNA trafficking, local translation, and stress.

## RESULTS

### KIF5A is an RNA binding protein that interacts with synaptic ribosomal protein mRNAs

KIF5A is the most preferentially and highly expressed kinesin in the brain based on gene expression data from the Genotype-Tissue Expression (GTEx) project^22^, (**Fig. 1a**), indicating its specialized functions in neurons. As neuron homeostasis depends heavily on mRNA transport for local translation, we explore the role of KIF5A in mRNA transport. While it is appreciated that mRNAs are transported in membrane-less RNP complexes by motor proteins, it remains unknown whether KIF5A binds directly to mRNA or indirectly via adaptor proteins^10,23–25^. Utilizing our Hydra deep-learning model^26^, we predict that KIF5A, as well as more than a third of human kinesin proteins, have RNA-binding capacity. (**Extended Data Fig. 1a**).

**Fig. 1.**
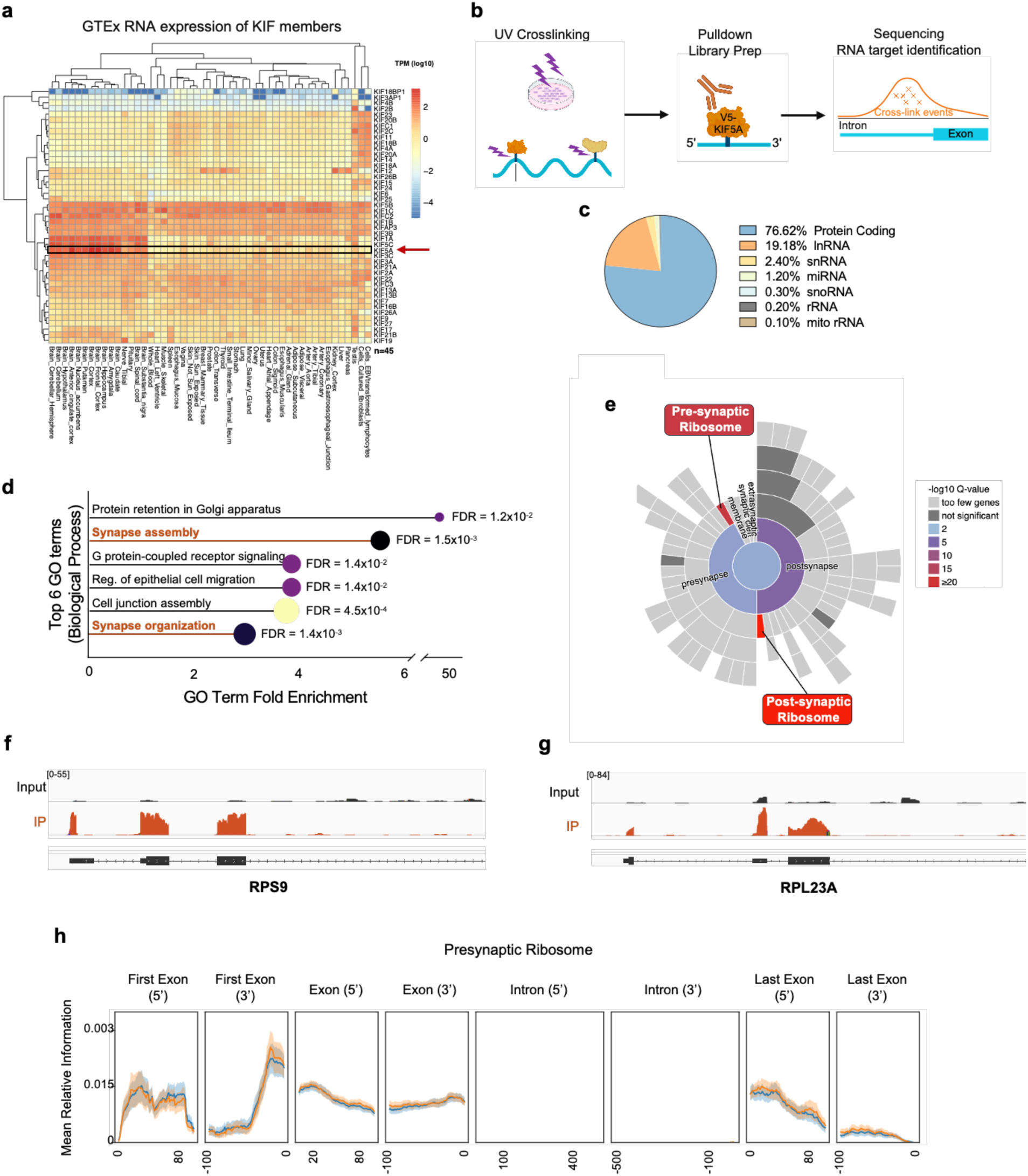
KIF5A is an RBP, binding to mRNA encoding ribosomal proteins. **(a)** Gene expression data of kinesins compiled from the GTEx project. **(b)** Schematic of eCLIP pipeline to identify KIF5A bound RNA. **(c)** Pie chart of genomic features represented in enriched windows in KIF5A eCLIP data from HEK293. **(d)** TOP GO terms for KIF5A RNA targets. **(e)** Statistically significant enriched SynGo terms for KIF5A RNA targets. **(f,g)** Significantly enriched KIF5A eCLIP-binding site on mRNA encoding RPS9 and RPL23a, respectively. **(h)** Mean relative information across pre-synaptic ribosome transcripts for KIF5A.

Focusing on KIF5A, we conducted enhanced cross-linking and immunoprecipitation (eCLIP)^27,28^ analysis on V5 tagged KIF5A in Hek293T cells (**Fig. 1b, Extended Data Fig. 1b,c**). Both eCLIP replicates passed statistical thresholds to maximize the number of hits (**Extended Data Fig. 1d**) and exhibited high concordance (**Extended Data Fig. 1e**) as specified by the Skipper computational analysis workflow^29^. After enforcing a 20% false discovery rate cutoff to define reproducible windows (**Extended Data Fig. 1f**), we observed that KIF5A binds primarily at coding sequences (CDS) and 5’ untranslated regions (5’ UTRs) within protein coding genes (**Fig. 1c**) (**Extended Data Fig. 1g**). Our results indicate that KIF5A interacts with mature mRNAs in the cytoplasm, resembling patterns of other RBPs that have CDS preferences in ENCODE3^30^.

We found that mRNA targets of KIF5A are statistically significantly enriched in terms that include “synapse assembly”, and “synapse organization” (**Fig. 1d**) when we carried out Gene Ontology (GO) analysis. Synaptic gene ontology (SynGO) analysis reaffirmed that KIF5A binds to mRNA encoding pre-synaptic and post-synaptic ribosomes (**Fig. 1e**). To illustrate, KIF5A binds to exonic regions of ribosomal protein S9 (RPS9) and ribosomal protein L23a (RPL23A) (**Fig. 1f,g**). RPS9 encodes a post-synaptic ribosomal protein^31,32^ and is a component of the small 40S ribosomal subunit. Mutations in RPS9 has been shown to alter synaptic composition and excitability^33^. RPL23a is a component of synaptic 60S ribosomal subunit. RPL23a expression is a potential biomarker for traumatic brain injury^34^. Averaging across all presynaptic ribosome mRNAs (**Fig. 1h**) and postsynaptic ribosome mRNAs (**Extended Data Fig. 1h**) using metadensity plots, we find that KIF5A exhibit exonic region binding, consistent with KIF5A’s localization in the cytoplasm. Our results indicate that KIF5A is an RNA binding protein that interact with mature mRNA encoding presynaptic and postsynaptic ribosomal proteins.

### KIF5A controls the localization of ribosomal proteins at the synapse

As KIF5A is classical known as a motor protein, we hypothesized that KIF5A transports its mRNA targets RPS9 and RPL23a to the synapse. We differentiated motor neurons from induced pluripotent stem cell (iPSC-motor neurons) via a combination of WNT activator and dual SMAD inhibitors^35,36^ (**Extended Data Fig. 2a**). Over ninety-five percent of the cells express motor neuron specific markers Islet1, SMI31 and βIII-Tubulin as measured by immunofluorescence (IF) indicating our differentiation was highly robust (**Extended Data Fig. 2b,c**). We depleted KIF5A in the iPSC derived MNs using short-hairpin RNAs (shRNAs). Compared to iPSC-motor neurons treated with scrambled control shRNA (ctrl shRNA), those treated with KIF5A-targeting shRNA shows 74% reduction in KIF5A expression by RT-qPCR (**Extended Data Fig. 2d**). Synaptic fractionation followed by RNA sequencing (RNA-seq) (**Fig. 2a**) revealed that reduced KIF5A expression leads to diminished synaptic localization of KIF5A-bound mRNAs that encode for synaptic ribosomal proteins (**Fig. 2b, Extended Data Fig 2e,f**). These findings suggest that KIF5A is responsible for the synaptic localization of its mRNA targets. To further determine whether decreased synaptic mRNA localization results in reduced synaptic protein levels, we assessed the synaptic presence of RPS9 and RPL23a using immunofluorescence (IF) in neurons treated with control or KIF5A-targeting shRNA. We found that the fraction of PSD95-positive synapses containing RPS9 decreases significantly (p < 0.05, two-tailed Student’s t-test) in neurons treated with KIF5A shRNA compared with those treated with ctrl shRNA (**Fig. 2c,d**). Similarly, KIF5A-targeting shRNA treated neurons showed reduction in RPL23a protein localization to the synapse (**Extended Data Fig. 2g**). Together, these results indicate that KIF5A controls the synaptic abundance of RPS9 and RPL23a mRNA and protein.

**Fig. 2.**
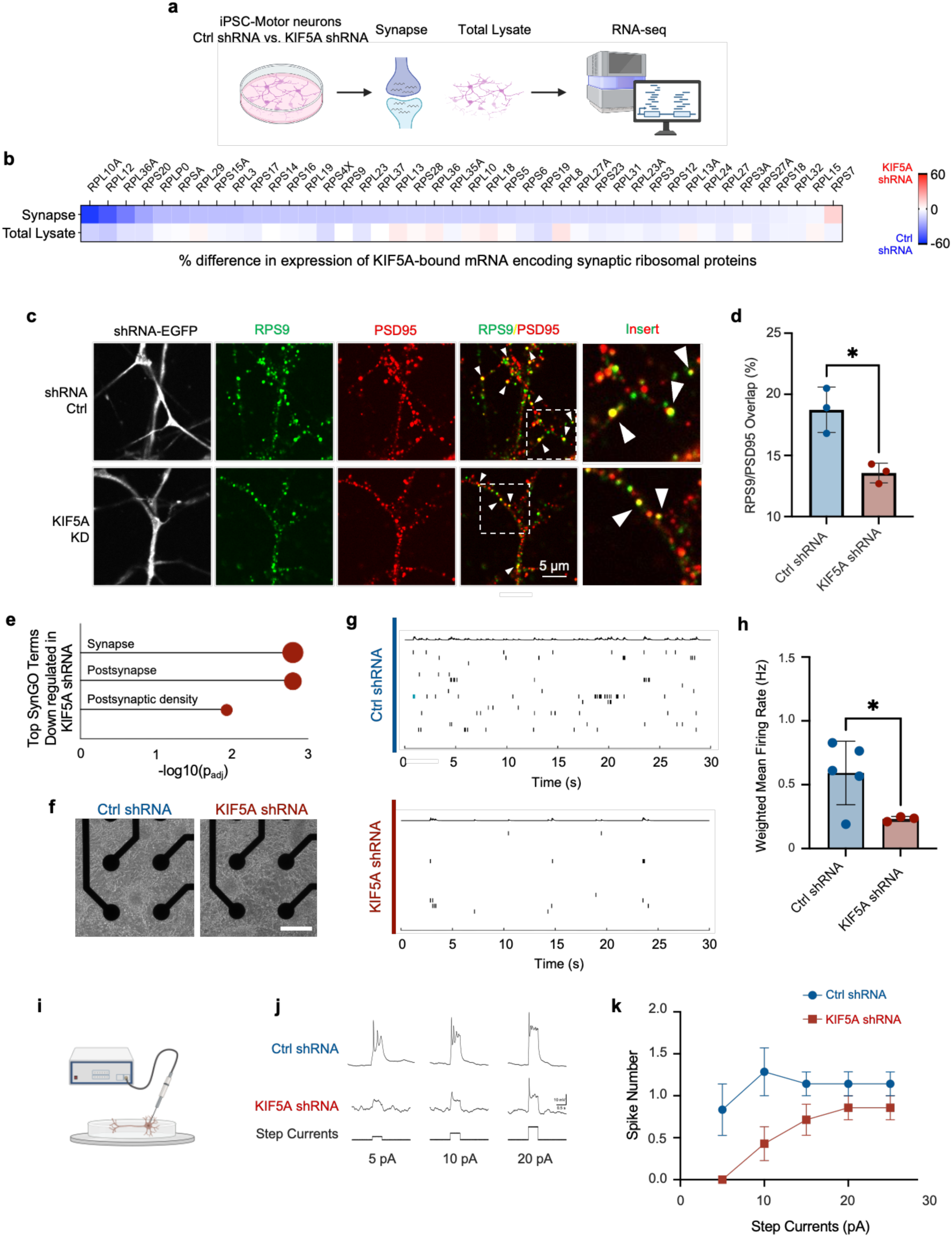
KIF5A is essential for RPS9 synaptic localization, synaptic composition, and function. **(a)** Schematic of synapse fractionation vs. total lysate for downstream RNA-seq analysis. **(b)** Heatmap showing percentage of difference in gene expression neurons treated with ctrl shRNA and KIF5A shRNA, in synapse and total lysate. Shown only synaptic ribosomal proteins whose mRNA bound by KIF5A. **(c)** Immunofluorescence co-staining of PSD95 and RPS9 in iPSC-derived motor neurons transduced with ctrl shRNA or KIF5A shRNA. shRNA lentiviral vectors carry EGFP as markers. Scale bars, 5 μm. **(d)** Percentage of PSD95, synapse marker, contains RPS9 from (a). N = 3, n > 300. *p < 0.05. Error bar = STD. **(e)** Statistically significant enriched SynGo terms for proteins that reduces in expression level in iPSC-derived motor neuron transduced with KIF5A shRNA compared to those transduced with ctrl shRNA, measured via proteomics. **(f)** Bright-field images of iPSC-derived neurons, treated with ctrl shRNA (left) or KIF5A shRNA (right), plated on MEA electrodes. Scale bar, 600 μm. **(g)** Examples action potential spikes in neurons treated with ctrl shRNA (top) or KIF5A shRNA (bottom). **(h)** Weighted mean firing rate of iPSC-derived motor neurons measured via MEA assay from (f). N > 3. *p < 0.05. Error bar = STD. **(i)** Schematic of patch-clamp experiment carried out in (i) and (j). **(j)** Example firing trace of individual iPSC-derived motor neurons carried out with current-clamp experiment and injected currents from 5pA to 20pA in 5pA steps. **(k)** Quantification of spike number measured in (h). N = 3, n = 11.

Synaptic ribosomal proteins such as RPS9 and RPL23a assemble into larger ribosomal subunits that control local translation of multiple synaptic proteins^37^. Since depleting KIF5A reduces the presence of RPS9 and RPL23a at the synapse (**Fig. 2c, Extended Data Fig. 2g**), we hypothesize that reduction in KIF5A expression may alter expression of other synaptic proteins. We performed proteomic analysis via label-free mass spectrometry (MS) on iPSC-motor neurons treated with either ctrl shRNA or KIF5A-targeting shRNA (**Extended Data Fig. 2h**). We observed that depleting KIF5A leads to reduction in protein levels of multiple synaptic proteins including SYN2, SNAP23, and CAMK1. SynGO analysis^38^ of proteins downregulated by KIF5A loss shows enrichment for postsynaptic proteins (**Fig. 2e**). Together, this indicates that KIF5A plays a broader role in maintaining synaptic protein levels and composition, exemplified by KIF5A controlling RPS9 and RPL23A expression.

### KIF5A is required for normal synaptic activity in human neurons

As depletion of KIF5A reduces ribosomal protein localization at the synapses which is likely to affect synapse protein composition and function^39–41^, we evaluated if electrophysiological properties of iPSC-motor neurons is modulated with KIF5A-targeting shRNA (**Fig. 2f,g**). An average weighted mean firing rate of 0.54 Hz was obtained by multi-electrode array (MEA) analysis on untreated iPSC-motor neurons (**Extended Data Fig. 2i**), consistent with previous studies ^42–44^. While ctrl shRNA did not alter the firing rates of these neurons (**Extended Data Fig. 2i)**, neurons treated with KIF5A-targeting shRNA exhibits a significant reduction in the firing rate (0.14 Hz; p < 0.05, two-tailed Student’s t test) (**Fig. 2h)**. We performed whole-cell patch clamp recordings on individual iPSC-motor neurons treated with ctrl shRNA or KIF5A shRNA (**Fig. 2i**). Neurons treated with ctrl and KIF5A shRNA exhibit similar resting membrane potentials of −33mV and −29.9mV respectively (**Extended Data Fig. 2j**), and thresholds of −23.6 mV and −22.9 mV respectively (**Extended Data Fig. 2k**), again consistent with previous studies^42,43^. To mimic physiological scenarios, we carried out current-clamp experiments and injected currents from 5 pA to 25 pA in 5 pA steps (**Fig. 2j**). We found that depletion of KIF5A reduces excitability of iPSC-motor neurons across 5 pA to 15 pA step currents (**Fig. 2k**), consistent with our MEA data. Together, our results indicate that KIF5A maintains electrophysiological homeostasis in human neurons.

### Mutant KIF5A neurons exhibit increased synaptic accumulation of RPS9 and RPL23a and neuronal activity

ALS-associated mutations in KIF5A are typically located in the C-terminal tail^15,16,20^, which is the cargo binding domain^44^ and can impact the movement of cellular cargo along neurites. Previous studies have demonstrated that multiple KIF5A ALS mutations cause exon 27 skipping, producing a truncated protein (KIF5AΔExon27) with a novel C-terminal tail^15,20,21^ (**Fig. 3a**). To understand how ALS-associated mutations in KIF5A alters neuronal function, we acquired fibroblasts from two ALS patients with mutations in exon 27 of KIF5A. One ALS patient carries a heterozygous substitution c.3019A>G and the other ALS patient has a heterozygous deletion c.2996delA (**Fig. 3b**). These fibroblasts were reprogrammed into iPSC (**Extended Data Fig. 3a**) and we confirmed these mutations via sequencing the genomic DNA (**Extended Data Fig. 3b**). We showed that c.3019A> G leads to exon 27 skipping (**Fig. 3c**) as measured by gel electrophoresis of RT-PCR products using RNA derived from patient iPSC-motor neurons. Exon 27 exclusion leads to truncation of WT protein and an addition of novel peptides (**Fig. 3d**), consistent with computational predictions in earlier reports^15,21^ The c.2996delA variant line, however, did not exhibit exon 27 skipping (**Fig. 3c**; **Extended Data Fig. 3c**). Interestingly, the deletion causes a frameshift and exclusion of the stop codon in exon 28 which consequentially leads to inclusion of the same novel peptides as the c3019A>G variant ^15,20,21^ (**Fig. 3d**).

**Fig. 3.**
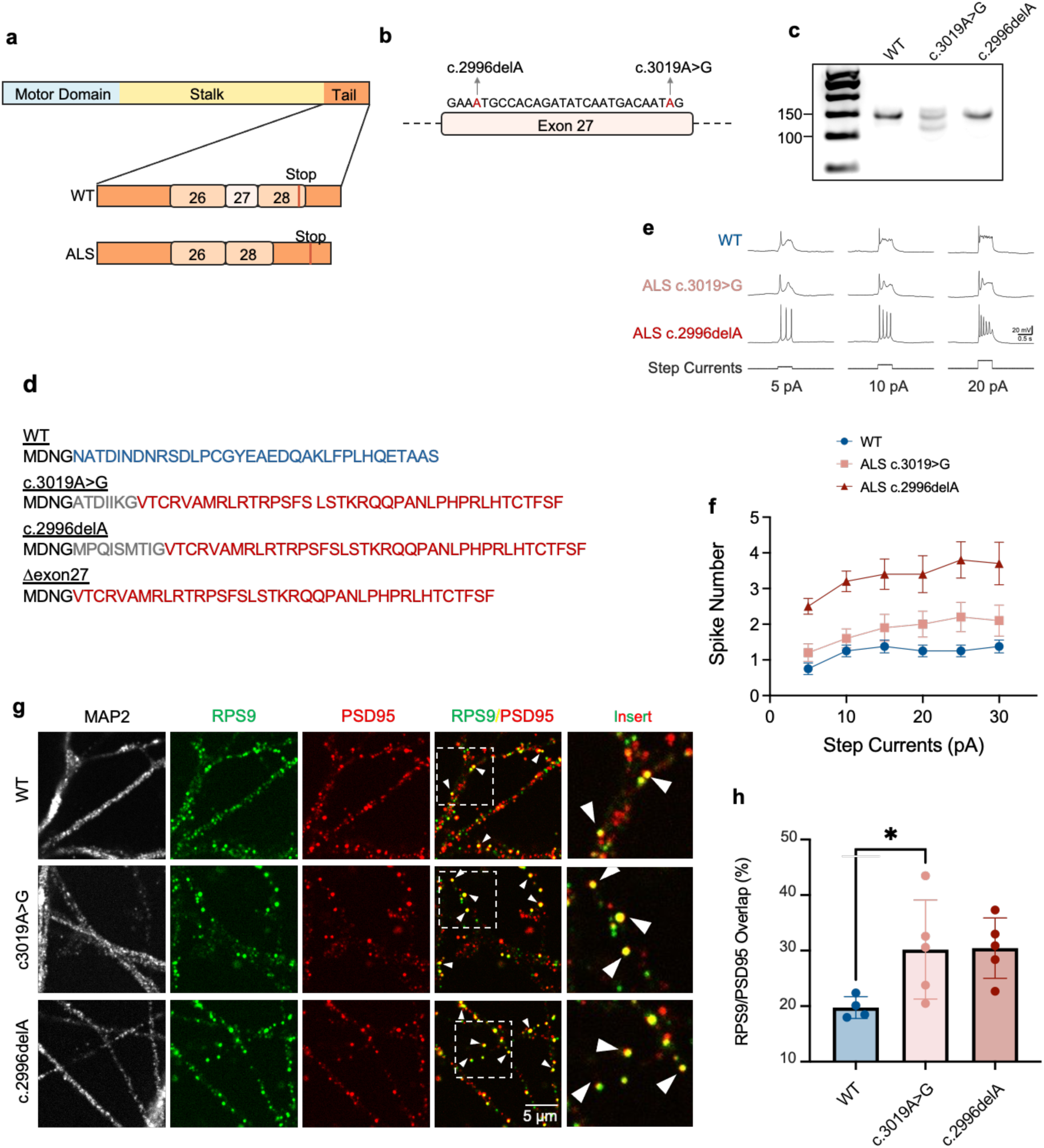
KIF5A ALS patient derived iPSC-motor neurons have abnormal function and synaptic composition. **(a)** Schematic of KIF5A domain structure and prediction that KIF5A ALS mutation leads to exon27 skipping and addition of novel peptide. **(b)** KIF5A ALS mutations in 2 patient lines used in this manuscript. **(c)** Gel electrophoresis of cDNA from exon 26 to 2xon28 to validate exon27 skipping in ALS patient lines. **(d)** ALS-associated mutations in KIF5A leads to a common C-tail as indicated in red. **(e)** Example firing trace of individual iPSC-motor neurons from healthy or ALS individuals. Experiments were carried out with current-clamp experiment and injected currents from 5pA to 20pA in 5pA steps. **(f)** Quantification of spike number measured in (e). N =3, n > 12. **(g)** Immunofluorescence of PSD95, RPS9 and MAP2 in iPSC-motor neurons from healthy and ALS individuals. Scale bars, 5 μm. **(h)** Percentage of PSD95, synapse marker, contains RPS9 from (g). N = 5, n > 300. *p < 0.05. Error bar = STD.

Having established that KIF5A is critical for synaptic function (**Fig. 2**) and recognizing that ALS-associated mutations in KIF5A typically occur in the cargo binding domain, we hypothesized that patient iPSC-motor neurons may exhibit altered firing properties. To test this, we performed electrophysiological recordings on patient-derived iPSC-motor neurons (**Fig. 3e**). Across all levels of step current stimulation, ALS patient neurons displayed higher firing rates compared to neurons from healthy individuals (**Fig. 3e and 3f**). Previous studies on electrophysiology of iPSC-motor neurons identified hyperexcitability in sporadic ALS (sALS)^45^, SOD1-^46^, and TDP43-mutant ALS^47^, and hypoactivity in FUS-mutant^48^, and c9orf72 ALS^49^. To our knowledge, this is the first observation that iPSC-motor neurons from KIF5A ALS patients exhibit hyperactivities resembling motor neurons from sALS, SOD1, and TDP43 patients.

We hypothesized that ALS-associated mutations in the cargo-binding domain of KIF5A might affect the synaptic localization of ribosomal proteins RPS9 and RPL23A. To test this, we performed immunofluorescence labeling for the synaptic marker PSD95 and quantified the levels of RPS9 and RPL23A at synapses in patient-derived and wild-type iPSC-motor neurons. We observed statistically significant (p<0.05, two-tailed Student’s t-test) increase of RPS9 accumulation at synapses in motor neurons derived from patients carrying the c.3019A>G and c.2996delA mutations compared to controls **(Fig. 3 g,h)**. Similarly, motor neurons from both ALS patients exhibited increased synaptic RPL23A levels relative to healthy controls **(Extended Data Fig. 3d)**. These findings indicate that motor neurons harboring KIF5A ALS mutations exhibit a phenotype opposite to that of KIF5A loss, indicating a gain-of-function effect that results in elevated ribosomal protein accumulation and synaptic hyperactivity.

### KIF5AExon27 expression increased synaptic localization of ribosomal proteins

ALS-associated variants c.3019A>G and c.2996delA both result in truncation of wild-type KIF5A and the addition of novel C-terminal peptide sequences **(Fig. 3d)**. We hypothesized that these structural changes confer gain-of-function properties, contributing to the observed hyperexcitability and increased accumulation of ribosomal proteins at synapses in ALS patient iPSC-motor neurons **(Fig. 3e-h)**. To test this, we overexpressed either wild-type KIF5A or a mutant form lacking exon 27 and containing the novel 39–amino acid sequence (KIF5AΔExon27)^15,21^ **(Fig. 3d)** in motor neurons derived from healthy individuals **(Fig. 4a)**. We confirmed that lentiviral delivery resulted in approximately a two-fold increase in KIF5A expression compared to non-transduced motor neurons, as intended **(Extended Data Fig. 4a,b)**. Importantly, both KIF5A WT and KIF5AΔExon27 constructs were expressed at comparable levels **(Extended Data Fig. 4c)**. Transcriptomic profiling revealed no significant differences between motor neurons expressing KIF5A WT and those expressing KIF5AΔExon27, suggesting that the mutant protein does not exert its effects via changes in steady-state mRNA levels.

**Fig. 4.**
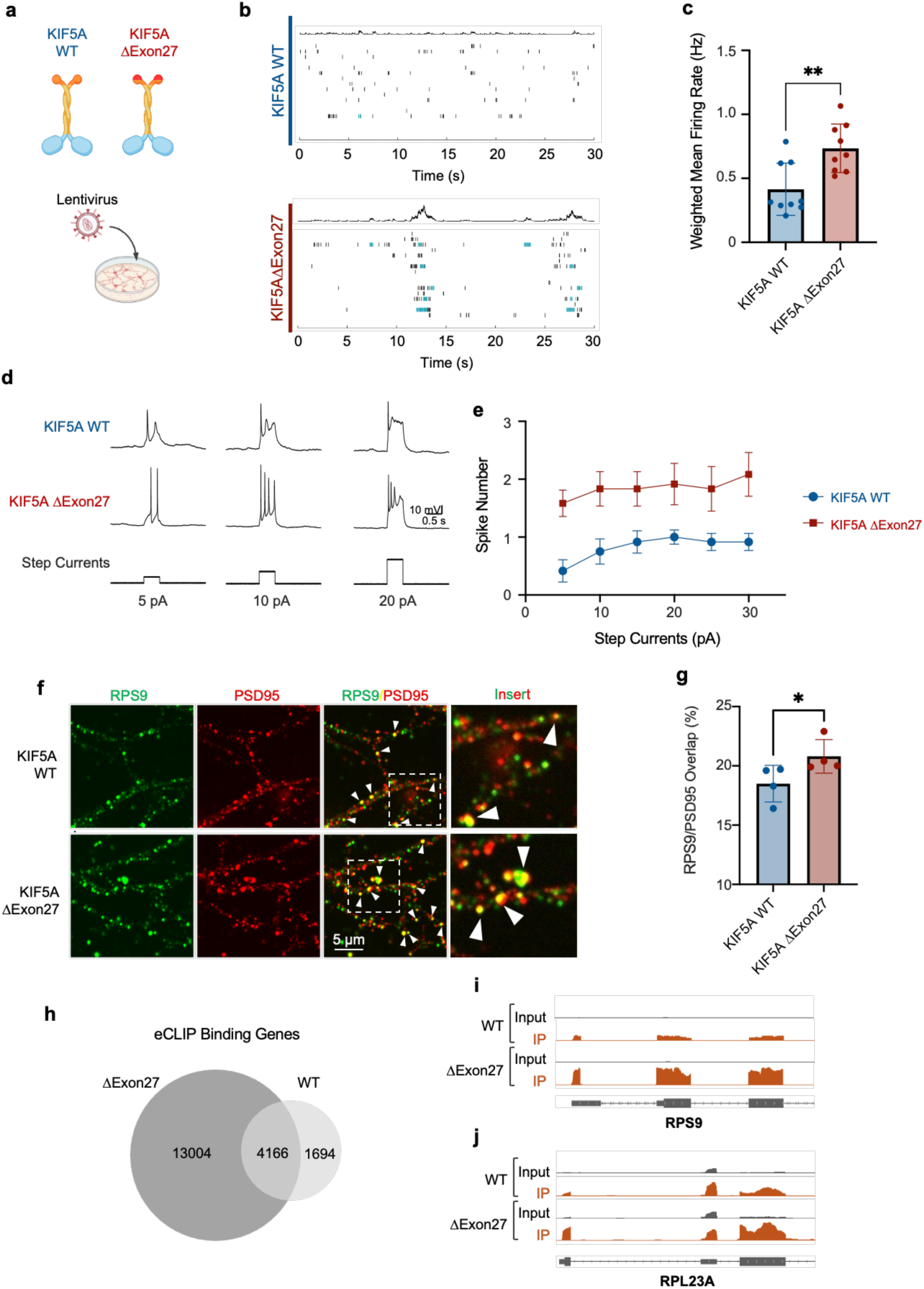
KIF5AΔexon27 is a gain-of-function, enhancing firing, synaptic localization of ribosomal proteins and enhanced mRNA binding. **(a)** Schematic of overexpression KIF5A WT or KIF5AΔExon27 in iPSC-motor neurons via lentivirus. **(b)** Examples action potential spikes in iPSC-motor neurons overexpressing KIF5A WT (top) or KIF5AΔExon27 (bottom). **(c)** Weighted mean firing rate of iPSC-motor neurons measured via MEA assay from (b). N = 9. **p < 0.01. Error bar = STD. **(d)** Example firing trace of individual iPSC-motor neurons carried out with current-clamp experiment and injected currents from 5pA to 20pA in 5pA steps. **(e)** Quantification of spike number measured in (D). N = 3, n ≥ 13. **(f)** Immunofluorescence of PSD95, RPS9 and MAP2 in iPSC-motor neurons overexpressed with KIF5A WT or KIF5AΔExon27. Scale bars, 5 μm. **(g)** Percentage of PSD95, synapse marker, contains RPS9 from (f). N = 4 n > 800. *p < 0.05. Error bar = STD. **(h)** Venn diagram of number of eCLIP binding genes overlapping between KIF5A WT and KIF5AΔExon27. Significantly enriched KIF5A eCLIP-binding site of KIF5A WT vs. KIF5AΔExon27 on mRNA encoding **(i)** RPS9 and **(j)** RPL23a.

To test whether KIF5AΔExon27 alters neuronal function, we conducted electrophysiological evaluations. Using MEA, we observed that motor neurons exogenously expressing KIF5AΔExon27 fires significantly (p < 0.01, two-tailed Student’s t-test) more frequently than those with KIF5A WT (**Fig. 4b** and **4c**). Our single-cell electrophysiological measurements using patch clamp analysis further supports our MEA findings (**Fig. 4d** and **4e**). This hyperactive firing pattern of motor neurons exogenously expressing KIF5AΔExon27 is similar to our findings in ALS patient-derived MNs with c.3019A>G and c.2996delA KIF5A mutations (**Fig. 3**), indicating that expression of KIF5AΔExon27 is the reason underlying motor neuron hyperactivity.

Given our observation that synaptic accumulation of ribosomal proteins is enhanced in motor neurons from patients carrying the c.3019A>G and c.2996delA KIF5A mutations **(Fig. 3g,h)**, we investigated whether exogenous expression of KIF5AΔExon27 would similarly alter the localization of these proteins. Immunofluorescence experiments revealed that expression of KIF5AΔExon27 in motor neurons led to increased synaptic localization of RPS9 compared to neurons expressing wild-type KIF5A **(Fig. 4f,g)**. Likewise, synaptic accumulation of RPL23A was also elevated in motor neurons overexpressing KIF5AΔExon27 **(Extended Data Fig. 4d)**. These findings support our conclusion that KIF5AΔExon27 promotes increased synaptic accumulation of RPS9 and RPL23A.

### KIF5AΔExon27 expression enhances binding to mRNAs

To determine whether the KIF5AΔExon27 mutant alters the RNA-binding properties of the protein, we performed eCLIP analysis in HEK293T cells expressing V5-tagged KIF5AΔExon27, following confirmation of successful immunoprecipitation (**Extended Data Fig. 4e**). Both replicates met stringent statistical thresholds and showed high concordance, as assessed using the Skipper computational pipeline **(Extended Data Fig. 4f-h**). KIF5AΔExon27 was found to predominantly bind protein-coding transcripts **(Extended Data Fig. 4i)**, with enrichment at coding sequences (CDS) and 5′ untranslated regions (5′ UTRs) **(Extended Data Fig. 4j)**, consistent with interactions with cytoplasmic mRNAs, similar to that of wild-type KIF5A.

Interestingly, KIF5AΔExon27 exhibited a higher proportion of binding to protein-coding mRNAs (97%) compared to wild-type KIF5A (76.62%) (**Extended Data Fig. 4i** and **Fig. 1c**) and bound a greater number of mRNA overall (**Fig. 4h**) despite comparable level of sequenced reads, indicating altered RNA-binding specificity or affinity. Notably, KIF5AΔExon27 showed increased binding to transcripts encoding RPS9 and RPL23A **(Fig. 4i** and **4j)**, which may explain the increased accumulation of these ribosomal proteins at synapses observed in motor neurons expressing KIF5AΔExon27, and by extension, in those carrying the c.3019A>G and c.2996delA mutations. Collectively, these findings suggest that KIF5AΔExon27 confers altered RNA-binding properties that could underlie its gain-of-function effects in ALS.

### Wild-type KIF5A interacts with RNA granule proteins and translation regulators

We next sought to investigate the protein environment surrounding KIF5A, with a particular focus on its association with ribonucleoprotein (RNP) complexes. To achieve this, we engineered a TurboID^50^ fusion at the C-terminus of KIF5A (KIF5A-TurboID) (**Fig. 5a**), incorporating a cleavable EGFP tag for fluorescence-activated cell sorting (FACS), which enables selection of TurboID-positive motor neuron progenitors for downstream assays (**Extended Data Fig. 5a,b**). In parallel, we generated a control construct tdTomato fused to TurboID (tdTomato-TurboID) to account for non-specific interactions, and a TurboID fusion with KIF5AΔExon27 (KIF5AΔExon27-TurboID) to model the ALS-associated truncation. Both constructs have a cleavable EGFP tag. Following lentiviral transduction and flow cytometry-based selection for comparable EGFP expression across all constructs (**Extended Data Fig. 5c-e**), we differentiated the selected clones into motor neurons and performed proximity labeling. Immunofluorescence confirmed robust biotinylation in neurites, validating the efficiency of our labeling approach (**Fig. 5b**).

**Fig. 5.**
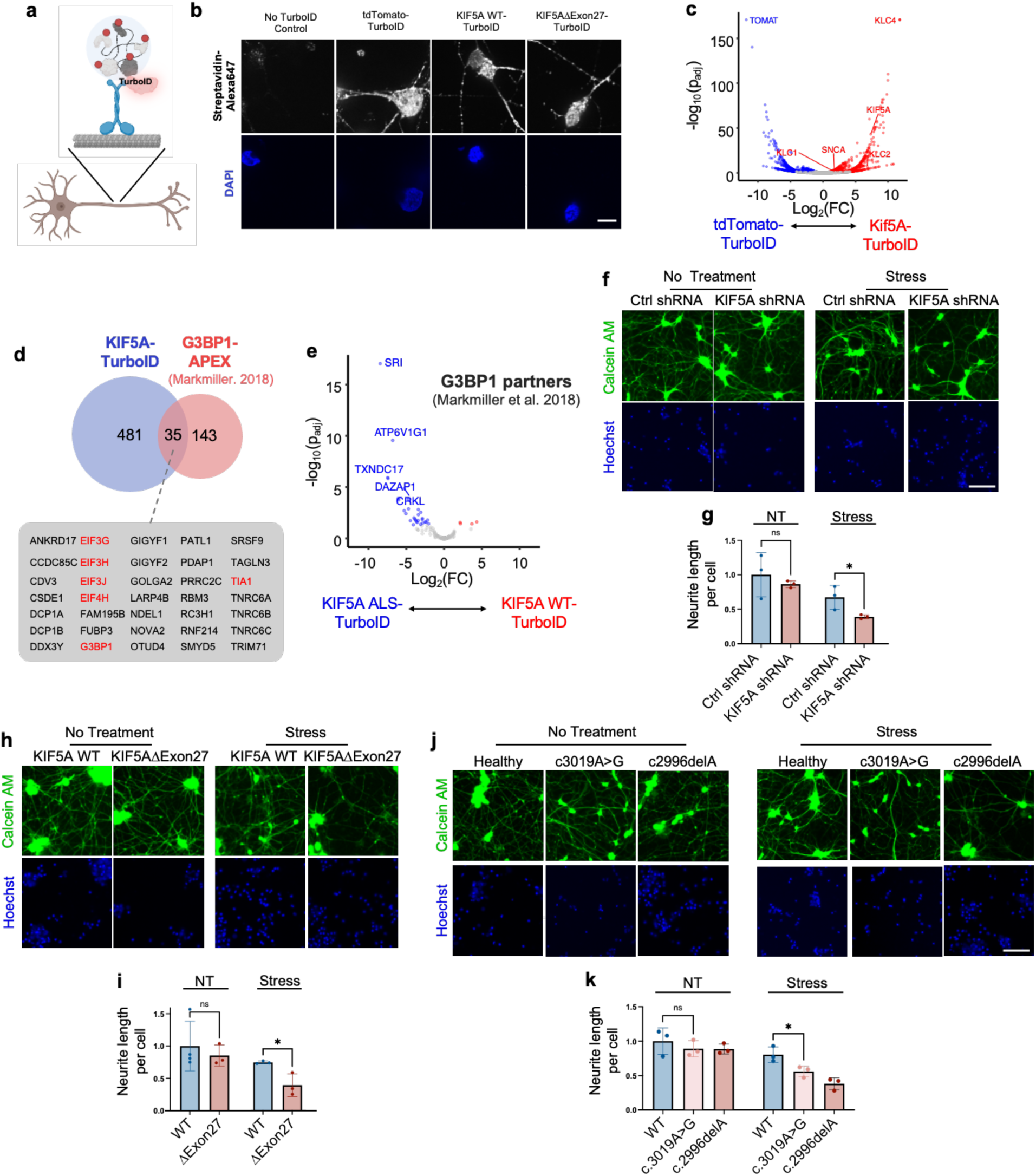
KIF5A WT interacting with G3BP1 partners; KIF5A ALS mutations enhanced binding to G3BP1 partners and susceptibility to stress. **(a)** Schematic of KIF5A-TurboID construct labeling proximal proteins. **(b)** Streptavidin staining of iPSC-motor neurons overexpressed with tdTomato-TurboID, KIF5AWT-TurboID or KIF5AΔExon27-TurboID. 500μm of exogenous biotin was added for 1 min. Scale bars, 10 μm.**(c)** Volcano plots showing statistically significant KIF5A-TurboID enriched prey proteins compared to tdTomato-TurboID control from proximity-labeling proteomics in iPSC-motor neurons. N = 3. **(d)** Venn diagram of KIF5A-TurboID hits versus G3BP1-APEX stress-granule hits. **(e)** Volcano plots showing statistically significant KIF5AΔExon27-TurboID enriched prey proteins compared to KIF5A-TurboID from proximity-labeling proteomics in iPSC-motor neurons. Only proteins overlapping with within G3BP1-APEX2 stress granule hits are shown. N = 3. **(f)** Staining of Calcein AM and Hoechst in iPSC-motor neurons treated with shRNA control or shRNA targeting KIF5A under non-treatment or puromycin treatment for 24 hours. Scale bar, 10 μm. **(g)** Quantification of neurite length per cell of neurons in (f). N = 3, n > 80. *p < 0.05. **(h)** Staining of Calcein AM and Hoechst in iPSC-motor neurons overexpressing KIF5A WT or KIF5AΔExon27 under non-treatment or puromycin treatment for 24 hours. Scale bar, 10 μm. **(i)** Quantification of neurite length per cell of neurons in (h). N = 3, n > 80. *p < 0.05. **(j)** Staining of Calcein AM and Hoechst in iPSC-motor neurons derived from healthy or ALS individials under non-treatment or puromycin treatment for 24 hours. Scale bar, 10 μm. **(k)** Quantification of neurite length per cell of neurons in (j). N = 3, n > 80. *p < 0.05.

Mass spectrometry analysis of the TurboID-labeled proteome revealed established KIF5A interactors, including KLC1, KLC2, KLC4, and SNCA^51^, confirming the specificity of our method (**Fig. 5c**). Interestingly, we discovered that KIF5A interacts with G3BP1 and a network of 35 G3BP1-associated proteins^52^, many of which are implicated in RNA granule biology and translational control (**Fig. 5d**). Among these, 20 proteins are directly involved in translation initiation (e.g., EIF3G, EIF3H, EIF3J, EIF4H, DDX3Y, FRUBP3)^53^, translation elongation (e.g., SRSF9, PDAP1, CSDE1, LARP4B), or translation repression (e.g., GIGYF1, GIGYF2, PATL1, TIA1, DCP1A/B, TNRC6A/B/C, TRIM71)^54,55^. These findings reinforce the emerging view that mRNAs are transported within neurites in a translationally repressed state^10,56,57^, and position KIF5A as a central scaffold linking RNA transport with machinery governing local translation.

### KIF5A’s role in stress response is compromised by ALS-associated alterations

Our data revealed that the ALS-associated KIF5AΔExon27 mutant alters RNA interactomes relative to KIF5A WT (**Fig. 4h-j**). As the mutations are in the cargo binding domain, we expect that KIF5AΔExon27 also alters protein interactomes. To test this, we carried out proximity labeling and mass spectrometry on motor neurons overexpressing KIF5AΔExon27-TurboID. We found that KIF5AΔExon27 interacts with a greater number of G3BP1 interactors than WT KIF5A (**Fig. 5e**), suggesting that the mutant protein perturbs the composition and dynamics of stress granules. Given the established role of G3BP1 in stress granule assembly^58,59^, we hypothesized that KIF5A may play an important role in neuronal stress granule biology. Furthermore, given the link between stress granule dysregulation and neurodegenerative diseases such as ALS^60,61^, we hypothesize that KIF5A mutations may disrupt the neuronal stress response pathway.

We depleted KIF5A in healthy iPSC-motor neurons and subjected the cells to puromycin-induced translational stress, subsequently assessing neuronal health via neurite tracing with Calcein AM dye (**Fig. 5f**). Under non-stress conditions, KIF5A shRNA-treated neurons exhibited neurite lengths comparable to those treated with ctrl shRNA; however, following puromycin treatment, KIF5A-deficient neurons displayed a pronounced reduction in neurite length, indicating heightened vulnerability to stress (**Fig. 5g**). Exogenous expression of KIF5AΔExon27 in motor neurons led to a similar decrease in resistance to translational stress, as evidenced by significant neurite retraction after puromycin exposure (**Fig. 5h,i)**. Importantly, motor neurons derived from ALS patients harboring KIF5A truncating mutations (c.3019A>G and c.2996delA), which mimic the KIF5AΔExon27 alteration, also exhibited diminished neurite outgrowth under stress conditions (**Fig. 5j,k**). Collectively, these results demonstrate that KIF5A is a previously unrecognized regulator of neuronal stress tolerance and that ALS-associated mutations compromise this protective function, thereby increasing neuronal susceptibility to stress-induced degeneration.

## DISCUSSION

Highly polarized neurons depend on active transport of RNA along microtubules to enable local protein synthesis at sites distant from the nucleus^10,24,56^. While it is widely appreciated that mRNAs are transported within RNP complexes by motor proteins^10,24,62,63^, the extent to which motor proteins directly interact with mRNA and whether distinct kinesin family members confer cargo-specific RNP composition remain largely unresolved.

Surprisingly, we discovered that KIF5A, a neuron-specific kinesin, functions as an RNA-binding protein. We then identified that KIF5A preferentially bind to mRNAs encoding synaptic ribosomal proteins such as RPS9 and RPL23a. Furthermore, we showed that KIF5A is crucial in governing the synaptic localization of these mRNAs and thus proteins, and proper synaptic activity of neurons. Beside binding directly to RNA, KIF5A also interacts with G3BP1 and its associated RNP granule components. Finally, we showed that ALS-associated KIF5A mutations alter mRNA and protein interactions, synaptic localization of ribosomal proteins, electrophysiology properties, and stress response of neurons. Based on our observations, we propose two mechanisms governing KIF5A role in ALS. The first features altered binding to mRNA encoding synaptic proteins, leading to synaptic dysfunction, and the second, altered proteins interactions with G3PB1 membrane-less RNA granules, leading to enhanced vulnerability to stress (**Fig. 6**).

**Fig. 6.**
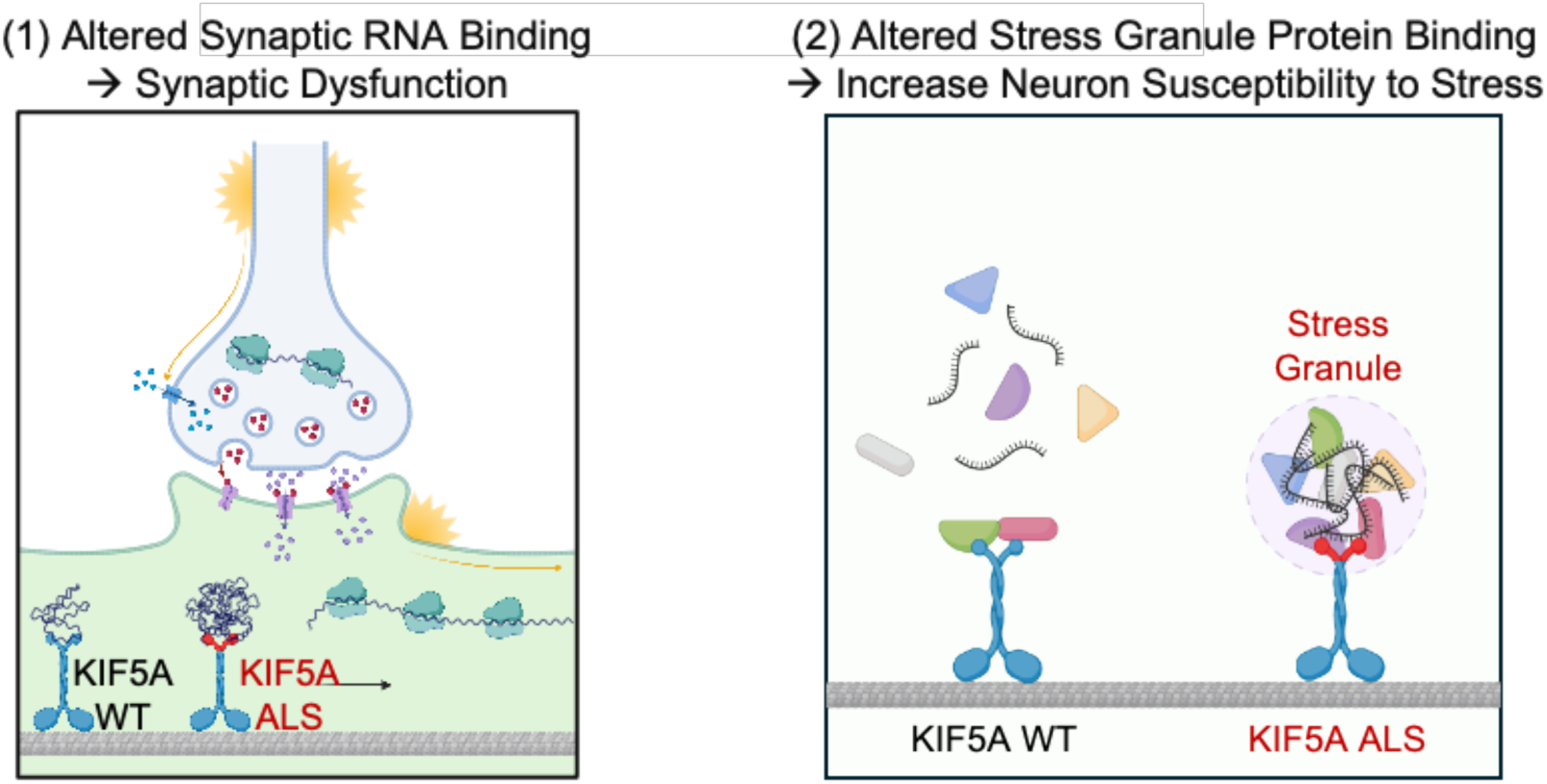
Schematic representation of mechanism of KIF5A dysregulation in ALS.

This study builds on decades of research into local translation in neurons, beginning with early electron microscopy studies in the 1970s that identified polyribosomes in dendritic spines^7,8^, suggesting that neurons transport mRNAs for local translation at distal synaptic sites. More recent work has confirmed the active transport of mRNAs in neurites^64–66^, but until now, the specificity of mRNA cargo and the mechanisms of their transport—particularly the direct involvement of motor proteins—remained unresolved.

Previous studies conducting high-throughput screens using poly-A capture^67^ and organic phase separation-based RBP discovery^67^ approaches suggested that several kinesins, including KIF1B, KIF1C, KIF2A, and KIF5B, could be RNA-binding proteins. However, these studies did not identify KIF5A as an RBP, likely due to their reliance on non-neuronal cell lines with low KIF5A expression. Our work provides the first computational and experimental evidence that KIF5A is indeed an RBP, preferentially binding mature mRNAs at coding sequences and 5’ UTRs, especially those encoding synaptic ribosomal proteins including RPS9 and RPL23a.

Functionally, we demonstrate that KIF5A is essential for the synaptic localization of RPS9 and RPL23a. This supports the centrality of mRNA transport for local translation at the synapse^1,37^. Furthermore, we demonstrate that KIF5A is essential for maintaining firing rates of neurons, reinforcing the connection between KIF5A, mRNA transport, local translation, and synaptic health. The clinical relevance of KIF5A is highlighted by its association with neurodegenerative diseases including ALS^15,20,21,68^, hereditary spastic paraplegia (SPG10)^69,70^, and Charcot-Marie-Tooth disease type 2 (CMT2)^18^. Notably, ALS-associated mutations in KIF5A cluster in the C-terminal cargo-binding domain, leading to exon27 skipping and truncated proteins with novel peptide sequences^15,20,21^. Previous studies have suggested that these mutations impair cargo binding and mitochondrial transport, resulting in hyperactive kinesin motors^20,21^. Our data expand these findings by showing that ALS mutations in KIF5A confer gain-of-function properties, including enhanced mRNA binding, increased synaptic accumulation of ribosomal proteins, neuronal hyperexcitability, and enhanced susceptibility to stress. These phenotypes are recapitulated in both patient-derived motor neurons carrying c.3019A>G mutation and c.2996delA mutation, and healthy neurons overexpressing the KIF5AExon27 mutant, demonstrating that the novel C-terminal peptide is sufficient to drive disease-relevant changes. Of note, we found that although c.2996delA mutation does not lead to exon27 skipping, the frameshift mutation also leads to truncation of KIF5A protein and addition of novel C-terminal peptide similarly to c.3019A>G mutation and other exon27 skipping KIF5A mutations reported previously^15,21^.

Finally, our study positions KIF5A within the broader landscape of mRNA transport, stress regulation and neurodegeneration^71^. We show that KIF5A interacts with RNA granule proteins such as G3BP1 and a network of translation initiation, elongation, and repression factors, supporting the model that mRNAs are transported in a translationally repressed state. G3BP1 plays a crucial role in stress granule formation; its dysfunction is implicated in multiple neurodegenerative diseases including ALS^72^. Here, we show KIF5AExon27 mutant exhibits increased binding to G3BP1 interactors. Both ALS patient-derived motor neurons and healthy neurons overexpressing the KIF5AExon27 mutant exhibit impaired stress responses and reduced neurite length following translational stress.

## CONCLUSION

This work uncovers a novel function for KIF5A as an RNA-binding protein that orchestrates the synaptic localization of ribosomal proteins and maintains neuronal homeostasis. Our data demonstrate that KIF5A is indispensable for proper synaptic composition and function, and that ALS-associated mutations in KIF5A confer gain-of-function properties that disrupt mRNA and protein interactions, leading to synaptic hyperexcitability and impaired stress responses. These mechanistic insights advance our understanding of the molecular underpinnings of ALS and highlight the centrality of mRNA transport, local translation, and stress response in neuronal health and disease. Targeting the aberrant mRNA trafficking pathways mediated by mutant KIF5A may represent a promising strategy for therapeutic development in ALS and related neurodegenerative disorders.

## METHODS

### Plasmids and Cloning

Plasmids were generated with PCR and NEBuilder Hifi cloning (ctrl shRNA, KIF5A shRNA, V5-KIF5A, and V-KIF5AΔExon27) or gateway recombination cloning (KIF5AWT-TurboID-P2A-EGFP, KIF5AΔExon27-TurboID-P2A-EGFP, and tdTomato-TurboID-P2A-EGFP). shRNA plasmids were generated with pLKO.1 puro vector (Addgene #8453), replacing PuroR with EGFP. Gateway cloning was carried out using pENTRY221 and pDONR221. Plasmids for lentivirus transduction in neurons were generated with pHR lentiviral vector.

### GTex analysis

KIF members expression was calculated by averaging RNA-seq TPM expression from The Genotype-Tissue Expression (PMID: 25954001). tissues with no expression in all of the KIF members were excluded. For clustered heatmap, infinite values (log10) were represented as −5.

### Lentiviral production

Lenti-X 293T cells (Takara Bio, 632180) were grown in high-glucose DMEM (Life, 11965118) supplemented with 10% fetal bovine serum (Life, 26140079) and subcultured using TrypLE Express (Life, 12604013) at a 1:10 ratio upon reaching approximately 90% confluence. For lentivirus productions, cells were seeded at 3.5×10^6^ cells per plate. After 24 hours, cells were cotransfected with the packaging plasmids psPAX2 (Addgene, 12260) and pVSV-G (Addgene #8454) together with in-house generated plasmids encoding either Ctrl shRNA, KIF5A shRNA, KIF5A WT, KIF5AΔExon27, or KIF5A-TurboID-P2A-EGFP, KIF5AΔExon27-TurboID-P2A-EGFP. Transfection was carried out using a Lipofectamine 3000. Following transfection, viral supernatants were harvested at 24, 48, and 72 hours, combined, and kept chilled. To concentrate viral particles, a Lenti-X concentrator (Takara Bio, 631232) solution was added, the mixture was incubated at 4 °C overnight, and then centrifuged at high speed and 4°C temperature. The resulting viral pellet was resuspended in phosphate-buffered saline at 1/200 of the initial supernatant volume. Viral concentration was determined with Lenti Go-Stix (Takaro Bio, 631280), and aliquots were frozen at −80 °C for storage.

### iPSC reprogramming and cell culture

iPSCs were reprogrammed from fibroblast via Sendai virus. iPSCs are maintained on 1× Matrigel (Corning, 354277) plates and mTSER Plus (Stem Cell Technologies, 100-0276)media. Media changes every 48 h. Cells were passaged using Versene (Life, 15040066) or Accutase (StemCell Tech, 07920) dissociation reagent at a 1:5–10 split ratio. For long-term storage, iPSCs are resuspended in a freezing solution containing 50% mTSER Plus media, 40% fetal bovine serum (FBS), 10% dimethyl sulfoxide (DMSO), and 10μM Rock Inhibitor (RI) (Tocris, 1254).

### iPSC-motor neurons differentiation

iPSCs are maintained on 1xMatrigel plate until they reach ∼90% confluency. On day 1-6 of motor neuron differentiation, cells are fed daily with N2B27 media with the addition of 1μM Dorsomorphin, 10 μM SB431542, and 3 μM CHIR99021. Day 7-18, cells are fed daily with N2B27 media with the addition of 1μM Dorsomorphin, 10 μM SB431542, 1.5 μM Retionic Acid (RA), 200 nM SAG and 5 μM RI. N2B27 media is made by adding 5mL N2 supplement (Invitrogen, 17502048), 10 mL B27 supplement (Invitrogen, 17504-044), 2mL of 50 mM ascorbic acid (Sigma, A4544), and 1% Pen/Strep to 500 mL of DMEM/F12 (Invitrogen, 10565042). On day 18, cells are at motor neuron progenitor (MNP) stage. During this MNP phase, cells are maintained and expanded in MNP media: N2B27 media supplemented with 3 μM CHIR99021, 2 μM DMH1, 2 μM SB431542, 0.1 μM RA, 0.5 μM Purmorphamine and 0.5mM valproic acid (VPA). For long-term storage, MNPs are resuspended in a freezing solution containing 50% MNP media, 40% fetal bovine serum (FBS), 10% dimethyl sulfoxide (DMSO), and 10μM Rock Inhibitor (RI) (Tocris, 1254). To further differentiate MNP to motor neurons, MNPs in motor neuron media are replated on PDL/PLO and laminin coated plates and counted as day 18 of differentiation. Motor neuron media is composed of 2 ng/mL GDNF, 2 ng/mL BDNF, and 2 ng/mL of GDNF in N2B27 media. On day 18 and d20, cells are fed with motor neuron media supplemented with 1.5 μM RA, 200nM SAG and 2 μM RI. On day 22 and d24, cells are fed with motor neuron media supplemented with 2 μM DAPT, 2 μg/mL laminin and 2 μM RI. Day 25 forward, cells are fed every 48 hr with motor neuron media supplemented with 2 μg/mL laminin and 5 μM RI (mature motor neuron media).

### MEA

On day 25, motor neurons are seeded on PDL/PLO/laminin coated 48-well plate CytoView MEA plate at 335,000 cells/well, using Accumax (Stemcell Technologies, 07921) as cell disassociating agent. On day 26, neurons are transduced with lentivirus of interest in 200 μL of mature motor neuron media. On d27, 300 μL of fresh media is added. Motor neurons are then fed every other day until d32. Before MEA experiments, 15 mM HEPES are added into the media via half media change. MEA experiments were carried out with Maestro Classic MEA system (Axion Biosytems)at the Human Embryonic Stem Cell Facility at the University of California, San Diego. MEA signals were sampled at 12.5kHz with 200 Hz high-pass and 3 kHz low pass filter. An adaptive spike detection threshold was set at 5.5 times of the standard deviation for each electrode with 1-s binning. Data acquisition for each condition were acquired for 5 min. Thresholded spike data (.spk files) were analyzed using Axion NeuralMetricTool software (v.2.1.5), evaluating weighted mean firing rate per well.

### Single-cell Electrophysiology

Day 25 iPSC-motor neurons are seeded on GG-12-laminin coverslips (Neuvitro Corporation, GG-12-Laminin) in 24 well plate at 335,000 cells/well. On day 26, neurons were transduced with lentivirus of interest in 200 μL of mature motor neuron media. On d27, 300 μL of media is added. Motor neurons are then fed every other day until d32. Coverslips were removed from the incubator on day 32 and were placed in Tyrode’s solution (in mM: 140 NaCl, 5 KCl, 2 CaCl2, 0.8 MgCl2, 10 HEPES and 10 Glucose, pH 7.4 adjusted with NaOH, 300 mOsm). Recording of neurons were obtained with an internal solution (in mM: 120 K-gluconate, 20 KCl, 4 Na2ATP, 0.3 Na2GTP, Na2 phosphocreatine, 0.1 EGTA and 10 HEPES, pH 7.3, 305 mOsm) in 4-6 MΩ glass pipettes. <u>Lentivirus transduced neurons were visually identified by expression of EGFP.</u> Data collected with the current-clamp method after the formation of giga seal. Recordings were made using Multiclamp 700B amplifier (Molecular Devices). Signals were filtered at 4 kHz and sampled at 10 kHz with a Digidata 1440A analog-digital interface (Molecular Devices). pClamp 10.7 software (Molecular Devices) was used to record and analyze data.

### Immunofluorescence

Day 25 iPSC-motor neurons are seeded on PDL/PLO/laminin coated 18 mm circular glass coverslips #1.5 (Neuvitro Corporation, GG-12-Laminin) in 12 well plate at 500,000 cells/well. On day 26, neurons were transduced with lentivirus of interest in 400 μL of mature motor neuron media. On d27, 600 μL of media is added. Motor neurons are then fed every other day until d32. Neurons are fixed with 4% Paraformaldehyde (PFA) (Electron Microscopy Sciences, 157-4-1L) for 1 hr at room temperature, quenched with 0.1M glycine (Fisher, G46-500) in PBS for 10 min at room temperature, and incubated in PBS until next step. Neurons are permeabilized and blocked with 0.1% Triton-X (Sigma, X100-500mL) with 5% goat serum (Sigma, G9023) for 30 min at room temperature. Cells are then incubated with primary antibodies diluted in blocking buffer (0.01% Triton-X with 5% goat serum) overnight at 4°C. Cells were washed 5 times with PBS and incubated with secondary antibodies diluted in blocking buffer for 1 hr at room temperature. Cells were then stained with DAPI (Thermo Fisher, 36971) and washed 5 times with PBS. Cell coverslips were then mounted with Prolong diamond antifade mountant (Thermo, P36965) and sealed with nail polish. Images were acquired with Nikon X1 Spinning Disk confocal equipped with a Plan Apo ʎ 60x NA 1.40 oil objective, 4 laser lines (405 nm, 488 nm, 561 nm, and 540 nm), Prime 95B sCMOS camera, emission filters (455/50 nm, 525/36 nm, 600/50 nm, and 705/72 nm).

### Image Analysis of PSD95 and RPS9/RPL23a colocalization

Immunofluorescence images were collected in .nd2 format are converted to .tiff then channels are split in ImageJ. Images are fed into in-house Cell Profiler to extract neurite masks using EGFP or MAP2 markers, nucleus detection via DAPI stain, and soma mask. These images are converted to 8-bit in Image J before put into stack, imported into MATLAB. A custom MATLAB script was used for spot detections of PSD95, RPS9 amd RPL23a. Spot count was performed using the Multiple Target Tracking (MTT) Algorithm based in MATLAB^73,74^. In the MTT algorithm, spot detection in images is performed by evaluating each 7 × 7 window in the image using a generalized likelihood ratio test to decide if a spot fits a point spread function, assuming Gaussian noise. Thus the analysis accounts for local background values, rather than global intensities, which is beneficial to account for the nonuniformity of autofluorescence across a cell. The algorithm also subtracts each spot and repeats the analysis until all spots are detected, which is beneficial when a high spatial density of spots is present. PSD95 and RPS9/RPL23a are definied to be co-localized if the spot center coordinates are within 10nm of each other.

### Neurite Tracing Experiment and Analysis

Day 25 iPSC-motor neurons are seeded on PDL/PLO/laminin coated, optic clear 96-well plate (Revvity Health Sciences, 502099831 at 20,000 cells/well. On day 26, neurons were transduced with lentivirus of interest in 50 μL of mature motor neuron media. On d27, 50 μL of media is added. Motor neurons are then fed every other day until d32. For stress challenge experiments, neurons are treated with 5μg/mL puromycin on day 31 for 24 hrs. On day 32, cells are labeled with Hoechst (Thermo Fisher, R37605) and Calcein AM (ThermoFisher, C3100) at manufacturer’s recommended concentrations in phenol red free DMEM /F12 (Thermo Fisher, 2104125) supplemented with 20mM HEPES. Cells are the incubated for 30 mins at 37°C, 5% CO2 incubator and imaged with Keyence BZ-X fluorescence microscope. Images are then exported to .tif files and analyzed with an in-house Cell Profiler workflow. Briefly, cell nucleus and soma are segmented using Hoechst and Calcein AM signals respectively. Soma was then masked from Calcein AM images. These images are then thresholded and applied morphological skeleton to segment out neurites. Neurite length per cell is calculated as morphological skeleton neurite length per image divided by number of soma detected in that image. Normalized neurite length per cell is normalized to average neurite length per cell of ctrl shRNA or WT control in each data set.

### Synaptic Fractionation

The isolation of synaptic fractions followed modifications based on established protocols^75^. First, neurons grown in 10cm plate were washed and incubated for 5 minutes on ice in 5 mL of hypotonic buffer (20 mM Tris-HCl, pH 7.5; 10 mM KCl; 1.5 mM MgCl₂; 5 mM EGTA; 1 mM EDTA) supplemented with a protease inhibitor cocktail (Millipore Sigma, #539134) and 1 mM DTT, all handled on ice. After incubation, neurons were gently loosened by swirling the plate and collected into a tube. The cells were pelleted by centrifugation at 200 × g for 3 minutes at 4°C. The pellet was gently resuspended in 1 mL of cold Syn-PER reagent (Thermo Fisher, #87793), supplemented with the same protease inhibitor cocktail, 1 mM DTT, and 800 U/mL SUPERase-In RNase inhibitor (Thermo Fisher #AM2694), followed by a 10-minute incubation on ice. To disrupt the cells, the suspension was pipetted up and down five times using a P1000 tip before being transferred to a glass Dounce homogenizer (Fisher, #K885300-0002) on ice. Homogenization was performed with approximately 24 strokes, after which cell lysis efficiency (most neurons lysed, nuclei intact) was confirmed under bright-field microscope. The homogenate was centrifuged for 10 minutes at 1,200 × g at 4°C. The resulting supernatant was transferred into a new top. To the supernatant, 20 units of Turbo DNase (Thermo Fisher, #AM2238) were added. This mixture underwent an additional centrifugation at 15,000 × g for 20 minutes at 4°C. The resulting pellet, enriched for synaptic components, was resuspended in 150 μL TRIzol and rapidly frozen for downstream RNA-extraction and analysis.

### RNA-seq

A minimum of 500,000 cells were collected and pelleted for RNA-seq analysis. Total RNA was extracted using RNeasy Plus Micro Kit (Qiagen, 74034) following the manufacturer’s protocol. For long-term storage prior to RNA extraction, cell pellets were kept in RLT Plus and b-mercaptoethanol at −20 °C. RNA-seq libraries were constructed using the Illumina Stranded mRNA Prep Kit (Illumina, 20040532). For sample barcoding, IDT for Illumina indices (Illumina, 20027213) were used, and sequencing was carried out on a NextSeq 2000 with compatible flow cells. Sequencing reads were evaluated with FastQC (v0.11.8), trimmed with cutadapt (v3.4), and aligned with STAR aligner (v2.7.6a) against the human genome (GRCh38). Gene-level quantification was performed with featureCounts from the subread (v2.0.6) package using GENCODE v38 annotation. Differential transcript expression was assessed using DESeq2 (v1.36.0). Data visualization for gene expression clustering and differential gene expression was performed in R (v4.2.1).

### Proteomics

Neurons grown in 6-well plates was washed 2 times with PBS, then scrapped, and lysed in RIPA buffer (Sigma, R0278). The suspension was sonicated for 400s (10s on/30 s off cycles) and cleared by centrifugation at 17,000xg for 15 min at 4°C. Samples are then submitted to the Sanford Burnham Prebys Proteomics core for total proteomics analysis.

### Generation of stable motor neuron progenitor cells expressing TurboID construct

Motor neuron progenitor cells were seeded in Matrigel coated 12-well plate at 600,000 cells/well. 24 hr post plating, cells were transduced with lentivirus of interest (KIF5AWT-TurboID-P2A-EGFP, KIF5AΔExon27-TurboID-P2A-EGFP, and tdTomato-TurboID-P2A-EGFP) in 200 μL of motor neuron progenitor media. Cells were fed every other day until day 7. On day 7, cells were FACS sorted based on EGFP signals compared to non-transduced cells. FACS was carried out with BD Aria II sorter at the Stem Cell core at the Sanford Consortium for Regenerative Medicine. EGFP positive cells were then replated and expanded.

### TurboID Experiments

Motor neuron progenitor cells expressing TurboID constructs were differentiated into motor neuron in 6-well plate. On day 30, motor neurons were treated with 200 μM biotin for 10 min at 37°C. Neurons were then washed 5 times with ice-cold PBS and dislodged. Cell pellets were then collected via centrifugation at 4°C, snap freezed in liquid nitrogen, and stored at −80°C. Frozen cell pellets were then shipped to the Sanford Burnham Prebys proteomic core for biotin AP-MS workflow and profile total proteome for biotinylated proteins.

### eCLIP

Enhanced crosslinking and immunoprecipitation (eCLIP) was performed largely according to established protocols with minor adaptations^76^. Cells were UV crosslinked at 254 nm (4000 μJ/cm²) to covalently bind RNA-protein complexes. For each replicate, approximately 20–30 million cells were harvested per sample. The crosslinked cells were lysed in eCLIP lysis buffer (50 mM Tris-HCl pH 7.4, 100 mM NaCl, 1% IGEPAL CA-630, 0.1% SDS, 0.5% sodium deoxycholate) supplemented with protease inhibitors. Lysates were briefly sonicated and partially digested with RNase I to generate RNA fragments. After clarification, RNA-binding proteins (RBPs) were immunoprecipitated with Dynabeads conjugated to antibody specific to the target RBP and incubated at 4 °C with rotation. Size-matched input (SMInput) controls were simultaneously generated using total cell lysates without immunoprecipitation.

After immunoprecipitation, samples underwent stringent washes using high-salt and low-salt buffers to reduce non-specific interactions. Bound RNAs were dephosphorylated with FastAP and T4 PNK, followed by ligation of a pre-adenylated 3′ RNA adapter. Protein–RNA complexes were resolved on 4–12% Bis-Tris polyacrylamide gels and transferred to nitrocellulose membranes. Regions corresponding to the expected size of RBP-RNA complexes were excised, treated with Proteinase K, and the RNA was purified. Reverse transcription was performed with Superscript III using indexed primers, followed by exonuclease cleanup and ligation of a 5′ DNA adaptor. The resulting cDNA libraries were PCR-amplified, size-selected, and purified prior to sequencing. Both IP and input libraries were submitted for Illumina sequencing, and downstream analysis was carried out using the Skipper pipeline to identify RBP binding sites and enriched sequence motifs.

### qRT-PCR

A minimum of 500,000 cells were collected and pelleted for RNA-seq analysis. Total RNA was extracted using RNeasy Plus Micro Kit (Qiagen, 74034) following the manufacturer’s protocol. For long-term storage prior to RNA extraction, cell pellets were kept in RLT Plus and b-mercaptoethanol at −20 °C. cDNA was generated with SuperScript III reverse transcriptase (Thermo Fisher, 18080-093). qPCR was then carried out with Power SYBR Green (Thermo Fisher, 4367659) using BioRad CFX Opus 384 Real-Time PCR System. GAPDH was used as normalization control.

### ALS patient gDNA and cDNA mutation validation

gDNA and RNA from patient iPSC-motor neurons were extracted using AllPrep DNA/RNA Kit (Qiagen, 80204). RNA was then reverse transcribed using SuperScript III into cDNA. Both gDNA and cDNA were PCR-amplified with specified primers. Amplified samples were separated using gel electrophoresis and imaged with Azure gel imager. For sequencing, appropriate band was excised and purified using Monarch DNA Gel Extration Kit (New England Biolabs, T1020). Samples were sequenced at Plasmidsaurus.

### Statistics and reproducibility

No statistical method was used to predetermine sample sizes, and experiments were generally not anonymized or randomized during analysis. Statistical analysis were carried out in Graphpad Prism. We employed the two-tailed Student’s t test as specified in the figure legends to assess statistical significance, with the following significance levels: *p< 0.05, **p< 0.01. For imaging data and blots, we performed each experiment at least in triplicate. Co-localization imaging experiments (for example, PSD95/RPS9 staining) were performed on at least 300 synapses for each replicate. Neurite tracing experiments were performed on at least 80 cells for each replicate. The figures shown in the manuscript are a representative sample of each experiment (for example, Figs. 2a, 3g, 4f, and 5f,h,j).

## ACKNOWLEDGEMENTS

We thank members of the Yeo Lab (A. Vu, G. Nguyen, F. Tan, and S. Blue) for their technical assistance. We also acknowledge the staff at the Nikon Imaging Center at UCSD for imaging support, the Institute for Genomic Medicine Core at UCSD for Illumina sequencing assistance, and the Sanford Burnham Prebys Proteomics Core for assistance with affinity pulldown proteomics. This work was supported by the following funding sources: the Schmidt Science Fellowship (to P. Le); the National Institutes of Health (NIH) Pathway to Independence Award (1K99NS140635 to P. Le); the Schmidt AI in Science Fellowship (to P. Le and N. Lal); the Branco Weiss Fellowship (to N. Lal); grant 2023-332369 from the Chan Zuckerberg Initiative DAF, an advised fund of Silicon Valley Community Foundation (to G.W.Y.); and the NIH (R01-HG004659 and R01-NS103172 to G.W.Y.). The funders had no role in study design, data collection and analysis, decision to publish, or preparation of the manuscript.

## AUTHOR CONTRIBUTIONS

P. Le and G. W. Y. conceived the projects. P. Le performed cell culture, sequencing, MEA, immunofluorescence, pulldown MS, eCLIP, synaptic fractionation, live-cell imaging experiments. N. Lal designed and performed TurboID experiments. N. Lal and D. Yang designed and performed single-cell electrophysiology. S. Xu analyzed RNA-seq and mass spectrometry data. S. Xu, B. Yee, H. Her and K. Rothamel analyzed eCLIP data. O. Mizrahi analyzed GTex data. M. Huang, S. Xu and S. Mumford assisted with molecular cloning, and cell cultures. Y. Mei assisted with synaptic fractionation experiments. B. Hoover and N. Schneider collected ALS patient fibroblast and reprogrammed them to iPSC. S. Blue assisted with eCLIP. G.W.Y supervised the project and manuscript preparation. P. Le wrote the initial draft of the manuscript, and all authors contributed to the final revisions.

---

**Extended Data Fig. 1.**
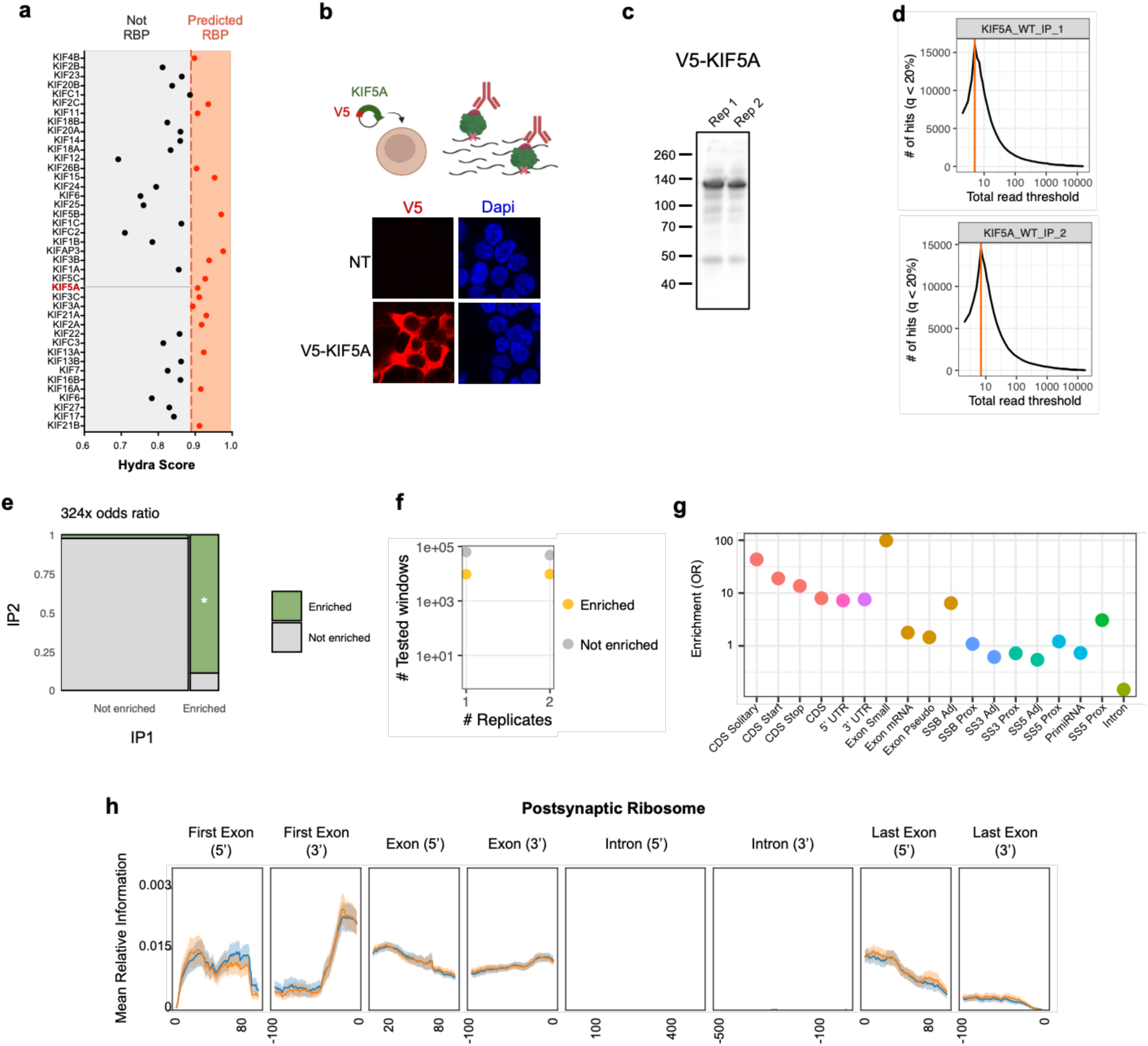
KIF5A quality control seCLIP analysis, related to Figure 1. **(a)** Machine learning prediction RBP identify of kinesins using Hydra. Proteins with Hydra score above 0.89 are considered RBP. **(b)** Top: Schematic of V5-KIF5A construct overexpressing in HEK293 cells and of cross-linking for seCLIP. Bottom: Immunofluorescence staining of V5 in cells non-transfected and transfected with V5-KIF5A. **(c)** Immunoblot of V5 after crossed linked and IP pull-down lysates from cells expressing V5-KIF5A. **(d)** Threshold scan produced by SKIPPER analysis of 2 IPs in iPSC-MNs. **(e)** Concordance between replicates. **(f)** Number of enriched windows and tested windows. **(g)** Enrichment odd ratio (OR) of seCLIP binding sites. **(h)** Mean relative information across post-synaptic ribosome transcripts for KIF5A.

**Extended Data Fig. 2.**
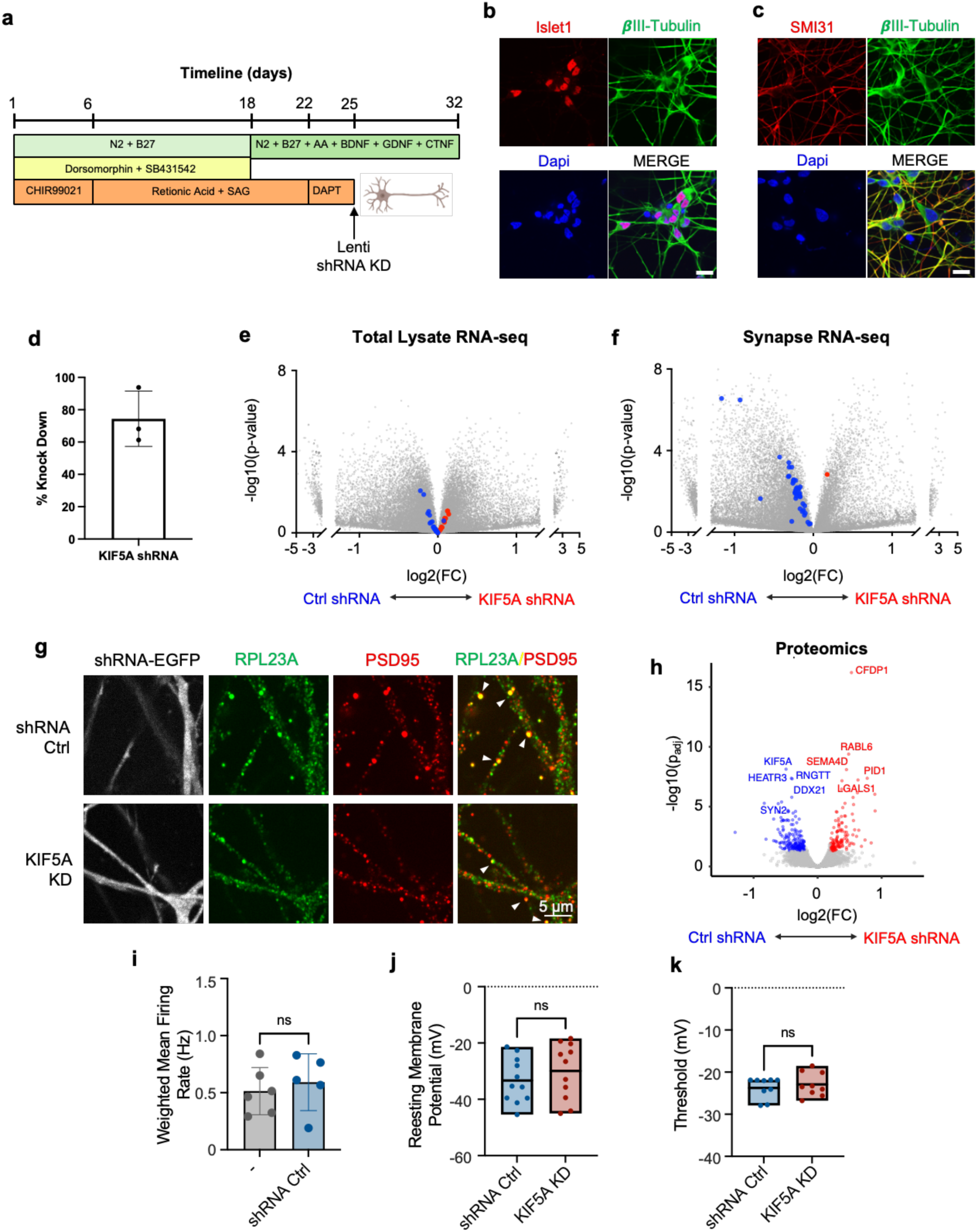
Motor neuron derivation from iPSC and characterization of iPSC-motor neurons with reduced KIF5A expression, related to Figure 2. **(a)** Schematic describing differentiation procedure from iPSC to motor neuron. For knockdown experiments with shRNA, lentivirus is transduced on day 25. **(b)** Validation of motor neurons markers Islet 1 and β-Tubulin via immunofluorescence. Scale bar, 20 μm. **(c)** Validation of motor neurons markers SMI31 and β-Tubulin via immunofluorescence. Scale bar, 20 μm. **(d)** Knockdown efficiency of shRNA targeting KIF5A in iPSC-MN measured by RT-qPCR. **(e,f)** Volcano plot showing differential gene expression of neurons treated with Ctrl shRNA versus KIF5A shRNA using (e) total lysate and (f) synaptic fraction. Blue and red dots represent synaptic ribosomal proteins whose mRNA bound by KIF5A. **(g)** Immunofluorescence co-staining of PSD95 and RPL23a in iPSC-derived motor neurons transduced with shRNA control or shRNA targeting KIF5A (KIF5A KD). shRNA lentiviral vectors carry EGFP as markers. Scale bars, 5 μm. **(h)** Proteomic analysis of iPSC-derived motor neurons treated transduced either shRNA control or shRNA targeting KIF5A. N = 3. **(i)** Weighted mean firing rate of iPSC-derived motor neurons non-transduced or transduced with shRNA control measured via MEA assay. N = 5. *p < 0.05. Error bar = STD. **(j)** Resting membrane potential of motor neurons transduced with shRNA control or shRNA targeting KIF5A, measured via patch-clamping individual neurons. N=3, n=11. **(k)** Threshold of motor neurons transduced with shRNA control or shRNA targeting KIF5A, measured via patch-clamping individual neurons. N=3, n=11.

**Extended Data Fig. 3.**
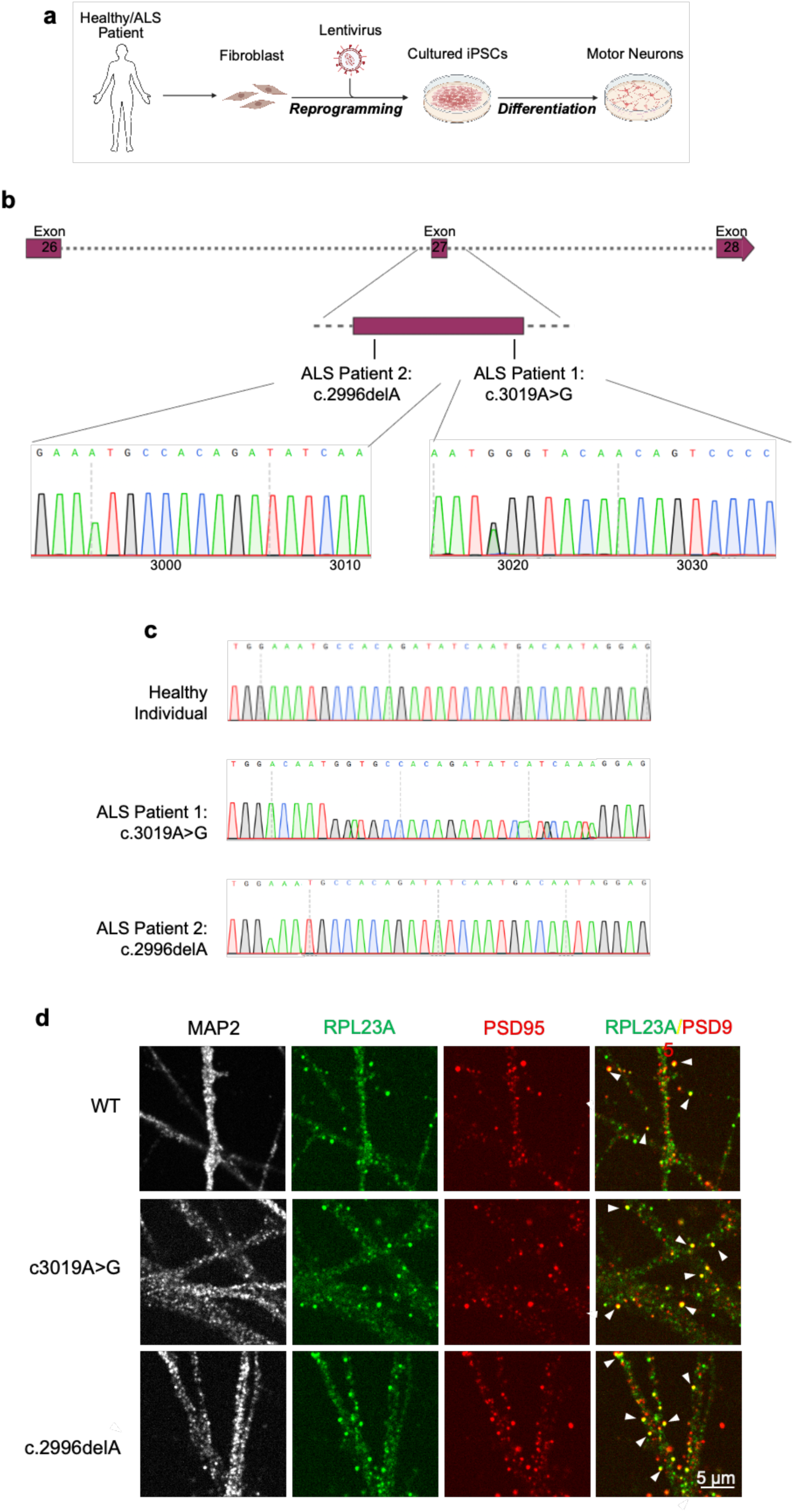
Characterization of ALS patient derived motor neuron, related to Figure 3. **(a)** Schematic of fibroblast reprogramming and iPSC-motor neurons differentiation. **(b)** Sequencing results of gDNA validating indicated mutations in ALS patient derived iPSC-motor neurons. **(c)** Sequencing results of cDNA validating indicated mutations in healthy and ALS patient derived iPSC-motor neurons. **(d)** Immunofluorescence co-staining of MAP2, PSD95 and RPL23a in iPSC-motor neurons from healthy and ALS individuals. Scale bars, 5 μm.

**Extended Data Fig. 4.**
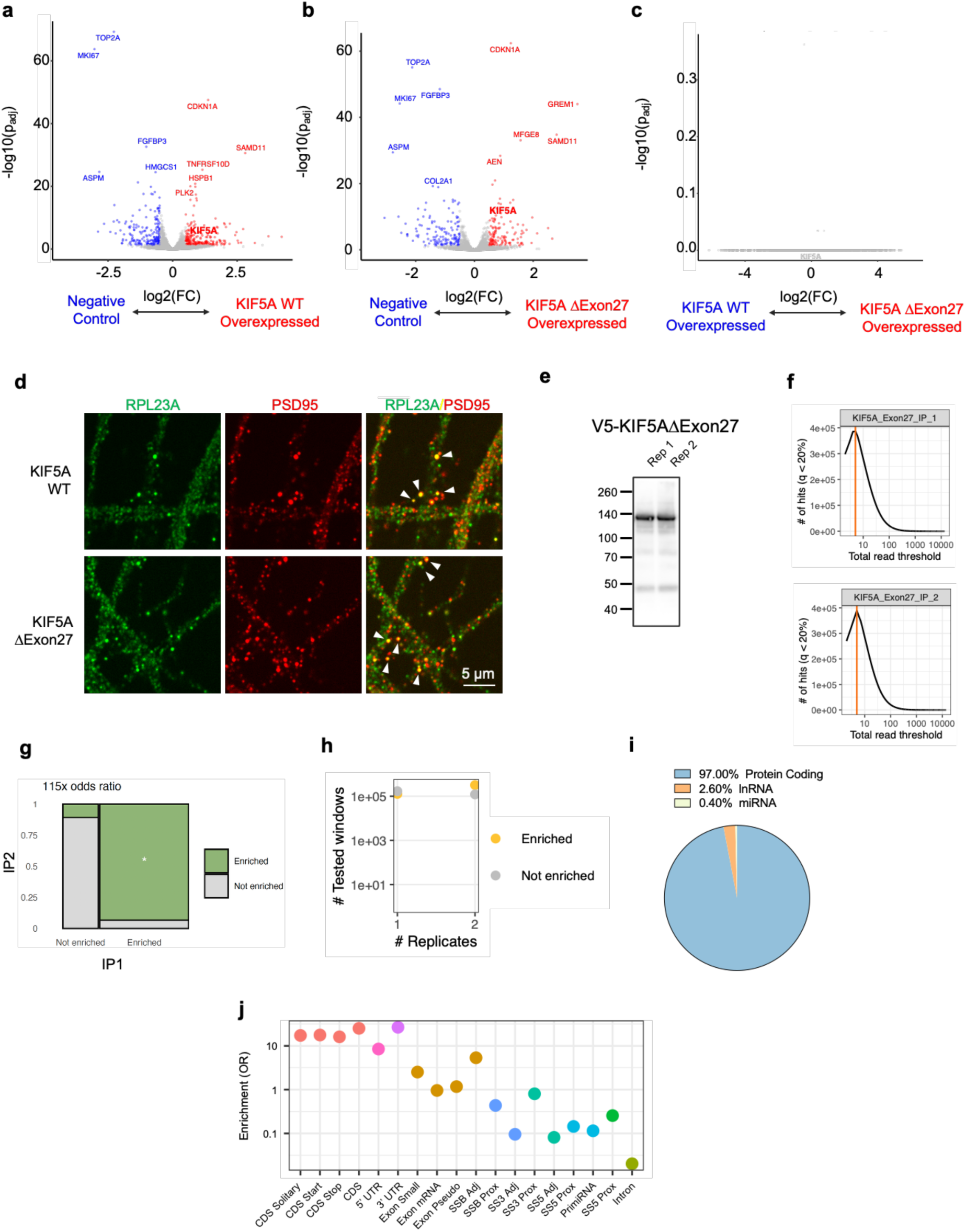
Characterization of iPSC-MN overexpressing KIF5A WT or KIF5AΔExon 27. KIF5AΔExon 27 quality control seCLIP analysis, related to Figure 4. **(a)** Volcano plot showing differential expression in iPSC-MN overexpressing KIF5A WT versus non-treatment control. **(b)** Volcano plot showing differential expression in iPSC-MN overexpressing KIF5AΔExon27 versus non-treatment control. **(c)** Volcano plot showing differential expression in iPSC-motor neurons overexpressing KIF5A WT versus KIF5AΔExon27 **(d)** Immunofluorescence co-staining of PSD95 and RPL23a in iPSC-motor neurons overexpressing KIF5A WT or KIF5AΔExon27. Scale bars, 5 μm. **(e)**Immunoblot of V5 after crossed linked and IP pull-down lysates from cells expressing V5-KIF5AΔExon 27. **(f)** Threshold scan produced by SKIPPER analysis of 2 IPs in iPSC-MNs. **(g)** Concordance between replicates. **(h)** Number of enriched windows and tested windows. **(i)** Pie chart of genomic features represented in enriched windows in KIF5AΔExon27 eCLIP data **(j)** Enrichment odd ratio (OR) of seCLIP binding sites.

**Extended Data Fig. 5.**
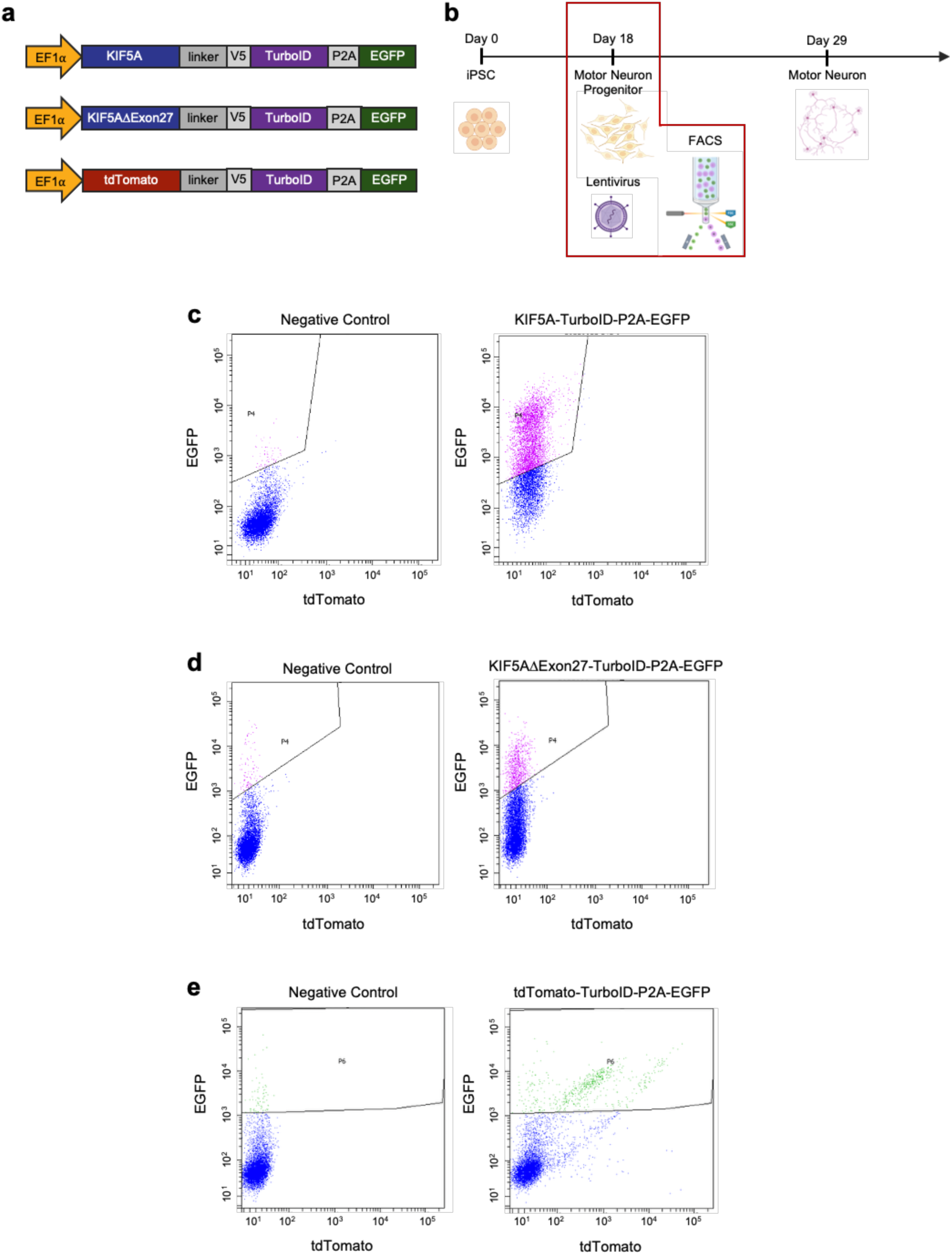
Generation of iPSC-MN expressing KIF5A-TurboID, KIF5AΔexon27-TurboID, and tdTomato-TurboID, related to Figure 5. **(a)** Schematic of plasmids encoding KIF5A WT-TurboID, KIF5AΔexon27-TurboID, and tdTomato-TurboID. **(b)** Schematic of generating stable motor neuron progenitors expressing plasmids in (a) via lentivirus. **(c)** FACS sorting based on EGFP expression for KIF5A-TurboID motor neurons progenitors. **(d)** FACS sorting based on EGFP expression for KIF5AΔexon27-TurboID motor neurons progenitors. **(e)**FACS sorting based on EGFP expression for tdTomato-TurboID motor neurons progenitors.

